# Helium as an *In Vitro* Furin Inhibitor

**DOI:** 10.1101/2020.12.18.423485

**Authors:** David Beihl

## Abstract

**Aims:** Certain cancers, pathogenic infections, and other diseases are facilitated by the host enzyme furin, a calcium-dependent serine protease that is the most prominent member of the family of proprotein convertases. Furin and the other proprotein convertases modify certain other proteins to change them from their inactive to active forms. Previous attempts to find an effective, non-toxic furin inhibitor to treat diseases facilitated by furin have had only limited success, due to toxicity or large molecular size that impedes absorption of the molecule. This has placed increased importance on the development of small-molecule furin inhibitors. The object of this study was to consider the effect of the noble gas helium as a furin inhibitor.

**Methods:** This study uses a fluorometric furin inhibition assay to compare the enzymatic activity of recombinant human furin after exposure to balloon-grade helium gas, compared to the enzymatic activity of untreated recombinant human furin.

**Results:** Helium exposure was found to decrease the *in vitro* enzymatic activity of recombinant human furin by as much as 55%; this inhibition lasted for up to 16 hours. Fluorescence measurements of enzymatic activity were taken for 24 hours.

**Conclusions:** These findings appear to be the first to report helium as a partial furin inhibitor. The observed partial inhibition was consistent throughout the experiment. The effectiveness of helium as a partial furin inhibitor, its favorable side-effect profile, and its long history of safe use in respiratory therapy, *when mixed with 20% oxygen or more and administered under direct medical supervision*, make it a promising treatment for diseases facilitated by furin or its substrates. Further studies in cell culture or clinical trials may expand its clinical role for such diseases.

## Introduction

The proprotein convertases are a group of enzymes, found in humans and other organisms, whose primary known function is to activate other proteins. In humans, there are presently nine recognized proprotein convertases. Each enzyme of this group operates by proteolytic cleavage of another protein’s inactive form (the proprotein), after which the protein’s activity is changed (usually increased). In some cases, the portion which is removed is also known to have biologic function. The proprotein convertases are generally felt to have overlapping, related but distinct functions from each other.

Viruses and bacteria are common causes of pathogenic infections in humans and other organisms. Many viruses rely on the proprotein convertases produced by their host to initiate and spread infection^1,2^. Certain bacteria also rely on these proprotein convertases, usually to activate bacterial toxins which cause disease^3,4^. In addition, a number of cancers, including malignant glioma^5^, astrocytoma^6^, cervical cancer^7^, and at least one colon cancer cell line^8^, also rely on furin or its substrates for malignant growth and/or metastasis. Therefore, medications that inhibit furin or other proprotein convertases might prevent or treat such diseases. However, the most commonly used proprotein convertase inhibitors have a narrow therapeutic index, which means the dose that inhibits the enzyme is similar to the dose that causes toxicity in the patient. One example of this is dec-RVKR-cmk. Others are complex biological products that can be difficult to synthesize or expensive to produce, or too large in size to enter cells or be absorbed effectively by the body or delivered to the affected organ or system. One example of this is alpha-1-antitrypsin-Portland^9,10^, a modified form of alpha-1-antitrypsin (which is about 44 kd in size). In addition, the efficacy and toxicity of many of these substances are often unknown, and information about the long-term side effects of their use is scarce if not absent. Thus, there is a clear and unmet need to develop effective small-molecule proprotein convertase inhibitors that can better prevent or treat bacterial, viral, neoplastic, or other diseases that are facilitated by proprotein convertase activity. The following experiment was conducted to evaluate the effectiveness of the noble gas helium as a furin inhibitor.

## Results

This experiment compared the activity of recombinant human furin with and without exposure to helium gas, using a microplate assay. The findings in Figures 1 and 2 demonstrate that helium caused up to 55% inhibition of furin activity; the inhibitory effect appears to have persisted for the first 16 hours during which the measurement took place. It should be noted, however, that estimates of the duration of this inhibition are not certain, because of apparent enzymatic degradation and substrate degeneration in the control wells late in the experiment, which can be seen in Figure 3. Also, the assay’s manufacturer has given no written guidance validating this assay for the purpose of testing the duration of furin inhibition beyond 1 hour; therefore, caution is advised regarding the exact duration of this effect. Furthermore, the results may underestimate the effect of helium inhibition because the experiment methodology erred on the side of allowing helium loss.

**Figure 1.**
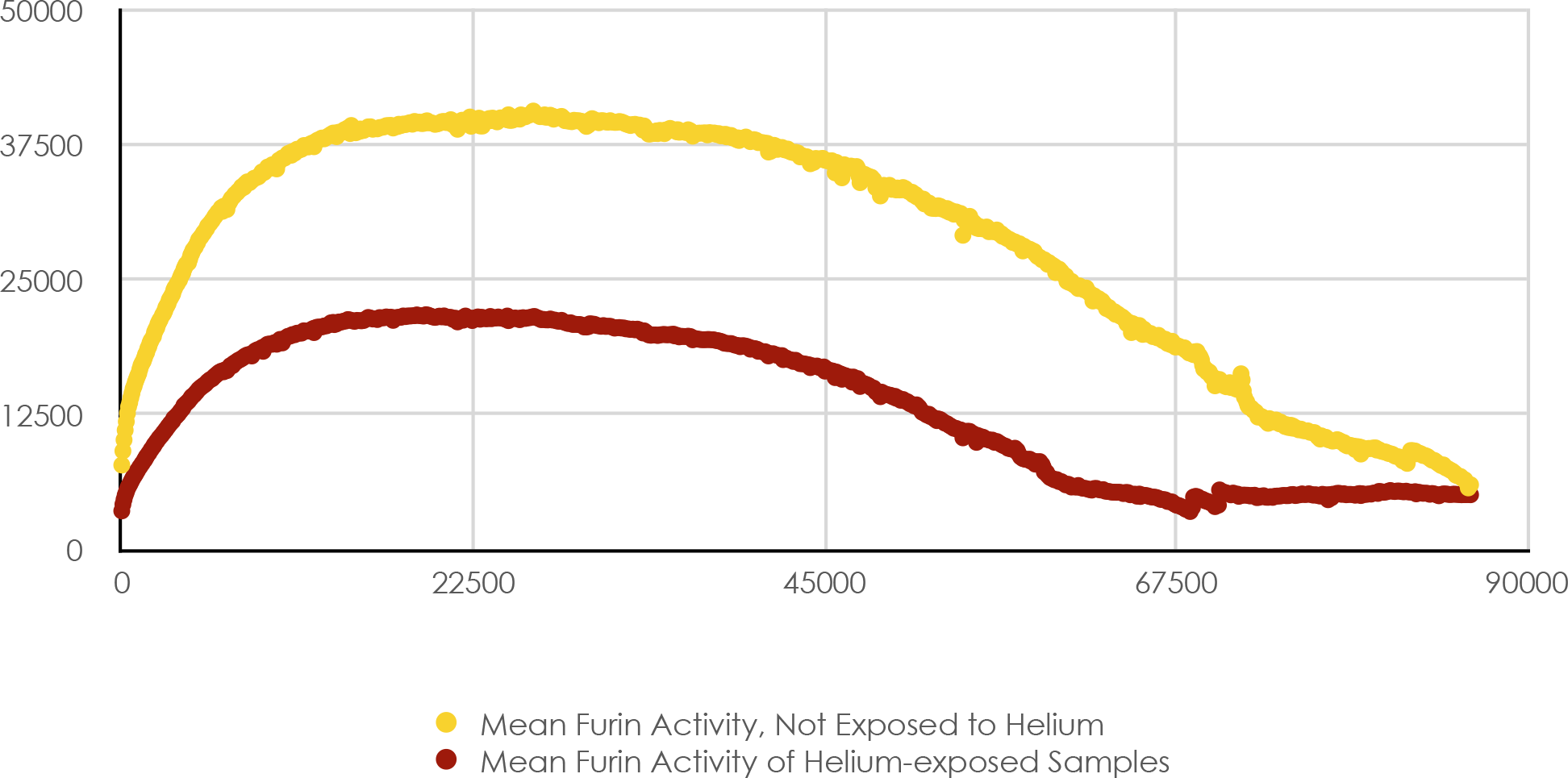
Mean Furin Activity (s-1) with and without Helium Exposure, Series E, vs Time (s)

**Figure 2.**
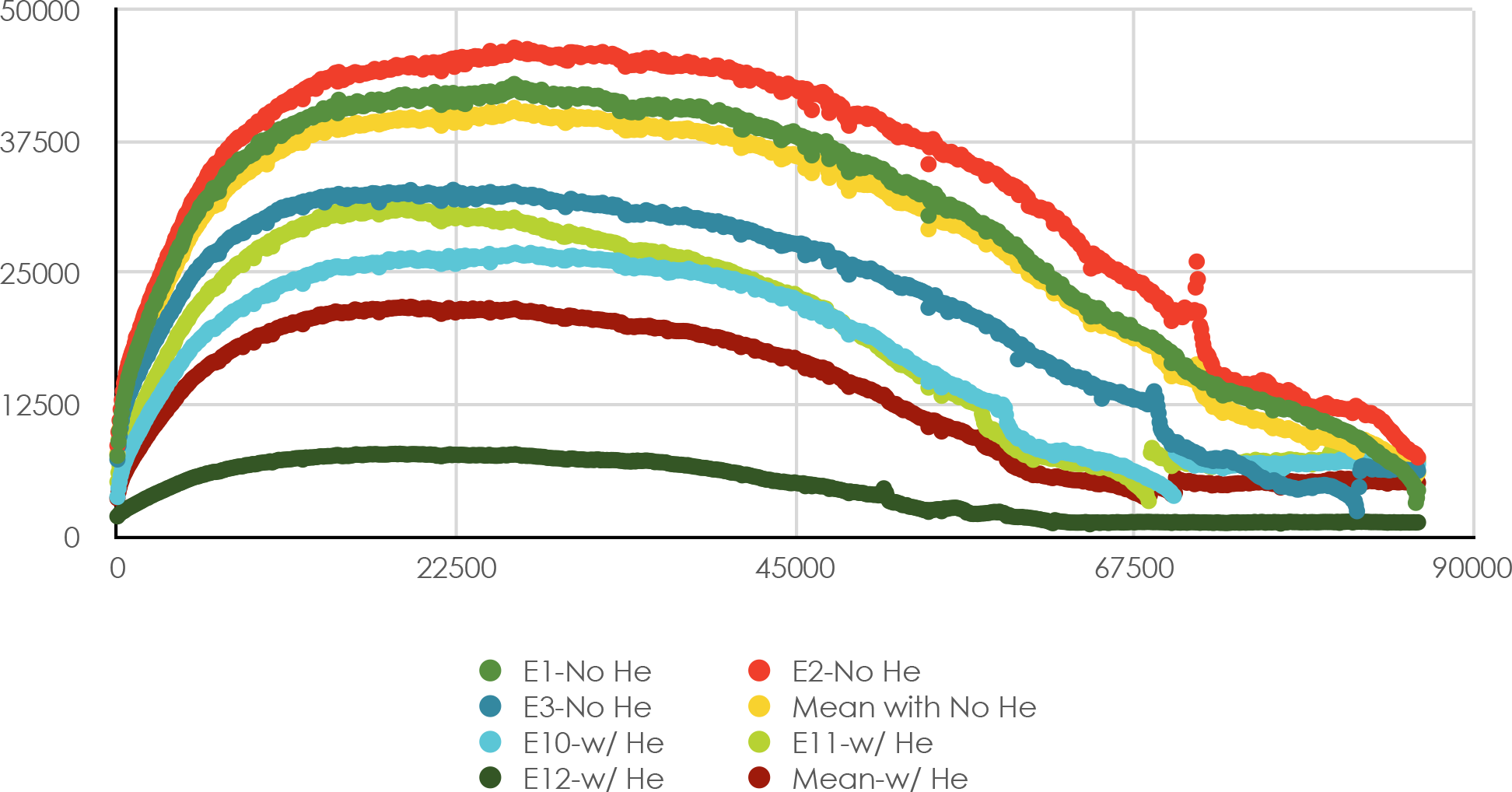
Mean Furin Activity (s-1) with and without Helium Exposure, Series E, vs Time (s)

**Figure 3.**
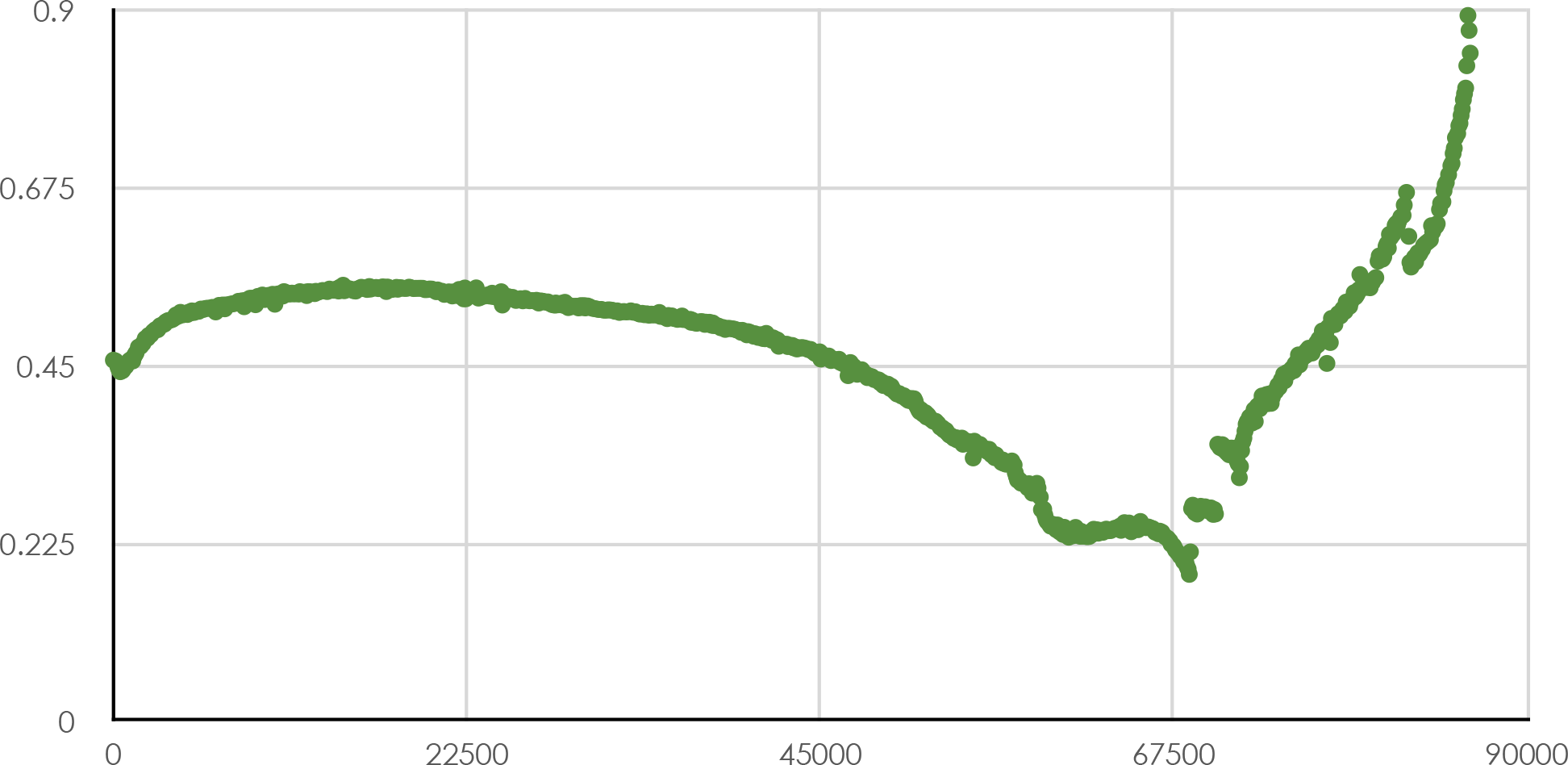
Mean Test/Control Furin Activity Ratio, Series E, vs Time (s)

The numbers on the vertical axis generally represent the amount of fluorescence measured in each well of the microplate during the experiment, which represents furin activity in this assay. The fluorescence tends to increase in the control wells over time because the uninhibited enzyme is actively cleaving the fluorogenic substrate, which continues to fluoresce after it has been cleaved and accumulates over the duration of the experiment, until it begins to degenerate. The wells exposed to helium exhibited decreased fluorescence as shown in Table 1 and Figures 1-3.

**Table 1.** Initial Subset of Data Used in Figures 1-3

Initial Subset of Data Used in Figures 1-3
|  |  |  |  |  |  |  |  |  |  |  |
| --- | --- | --- | --- | --- | --- | --- | --- | --- | --- | --- |
| *Time (s)* | 0.0 | 72.5 | 144.9 | 217.4 | 289.8 | 362.2 | 434.6 | 507.1 | 579.5 | 651.9 |
| E1-no helium (s<sup>-1</sup>) | 7620 | 8945 | 9978 | 10885 | 11694 | 12537 | 13150 | 13600 | 14072 | 14525 |
| E2-no helium (s<sup>-1</sup>) | 8584 | 9867 | 10978 | 12015 | 12905 | 13744 | 14505 | 14916 | 15467 | 15906 |
| E3-no helium (s<sup>-1</sup>) | 7269 | 8569 | 9514 | 10271 | 10959 | 11585 | 12043 | 12308 | 12654 | 12991 |
| Mean-no helium (s<sup>-1</sup>) | **7824** | **9127** | **10157** | **11057** | **11853** | **12622** | **13233** | **13608** | **14064** | **14474** |
| E10-w/ helium (s<sup>-1</sup>) | 3708 | 4528 | 5171 | 5749 | 6148 | 6557 | 6895 | 7094 | 7368 | 7649 |
| E11-w/ helium (s<sup>-1</sup>) | 5145 | 6014 | 6611 | 7138 | 7511 | 7914 | 8280 | 8515 | 8800 | 9093 |
| E12-w/ helium (s<sup>-1</sup>) | 1868 | 2007 | 2100 | 2210 | 2279 | 2363 | 2375 | 2479 | 2543 | 2620 |
| Mean-w/ helium (s<sup>-1</sup>) | **3574** | **4183** | **4627** | **5032** | **5313** | **5611** | **5850** | **6029** | **6237** | **6454** |
| Ratio of mean activity with helium/<br>mean activity with no helium exposure | **0.457** | **0.458** | **0.456** | **0.455** | **0.448** | **0.445** | **0.442** | **0.443** | **0.443** | **0.446** |

Figure 1 compares the mean furin activity with and without helium exposure from the final three control wells and three test wells of the microplate (the “E” wells, or “series E”). Of the samples available, these appear to be most representative of the inhibitory capability of helium on furin. A more detailed discussion of this matter is in the “Materials and Methods” section below, as well as in Appendix A.

## Discussion

This study appears to be the first to report helium’s effect to inhibit furin activity. The observed partial inhibition appears to have lasted for up to the first 16 hours of the experiment, and may have been limited by helium’s poor solubility in aqueous solution. It is possible that helium might even have a greater furin inhibitory effect *in vivo*, because the large surface area of the lungs promotes absorption and facilitates the access of helium to alveolar furin and cell membranes.

Of note, helium’s furin inhibitor effect explains some apparently unexplained previous findings of other researchers. According to one past report, pre-administration of 75% helium in an animal model of ischemic stroke inhibited tissue plasminogen activator-induced thrombolysis^11^. Notably, TPA is a furin substrate^12^, and its amino acid nsequence contains a furin cleavage site^13^. This raises the question whether the observed anti-TPA effect could have been mediated by helium inhibition of furin.

In addition, other investigators have reported using heated heliox (a helium-oxygen mixture) for human viral infections such as SARS-CoV-2. These researchers report early promising results with “dozens” (the exact number was not specified) of moderately ill COVID patients with lung lesions of 50-80%, indicating that all patients treated with heated heliox recovered, and that their COVID PCR tests generally became negative after 2-3 days. These authors seem to primarily have considered heliox as a delivery vehicle to transfer heated gas to the airway, and did not seem aware of helium’s furin inhibitory effect^14^. A third paper described the ability of heliox to decrease pulmonary inflammation, which was thought to be secondary to its ease of respiration^15^, rather than enzymatic inhibition. The novel finding of a furin inhibitory effect for helium may better explain these previous findings.

Given the novelty and non-obvious nature of this finding—a proprotein convertase inhibited by a noble gas—it is natural to inquire regarding its potential mechanism of action. One possibility is that helium could use a mechanism of action similar to whatever xenon’s mechanism of inhibiting furin is.^16^. Although xenon’s mechanism of furin inhibition is also uncertain, it may occur through Van der Waals interactions between the loosely attached outer valence electrons of xenon and the loosely bound electrons in the aromatic side chains of amino acid residues such as phenylalanine. One previous study has already shown that xenon’s widely recognized inhibition of neuronal NMDA receptors is mediated by phenylalanine interactions, and that mutating a crucial phenylalanine residue to tyrosine or tryptophan prevents xenon from inhibiting the NMDA receptor^17^. Another study has shown that furin has a highly hydrophobic pocket in the P domain, which is adjacent to the active site.^18^ This region contains phenylalanine at position 565, as well as tyrosine at position 560^19^. Under these circumstances, it is possible that xenon’s (and perhaps helium’s) inhibitory effect on furin is mediated by Van der Waals interactions with the aromatic side chains of Phe565 and Tyr560 in furin. This hypothesis could be tested by creating mutant forms of furin that change Phe565 to Tyr or Trp, as was done in the NMDA receptor study^20^, or by changing one or both of these to non-aromatic amino acids, and testing the modified furin’s degree of inhibition by xenon or helium. It is also possible that helium may bind elsewhere, causing unfavorable conformation change, or may use a different, unknown mechanism. X-ray crystallography and related studies may clarify this in the future.

Overall, the favorable side-effect profile of helium, and its long history of safety *when mixed with at least 20% oxygen and administered under direct medical supervision*, make it a promising treatment for the viral, bacterial, neoplastic, and other diseases facilitated by furin or its substrates, when no better treatment (such as xenon^21^) is available, approved by regulators, or clinically appropriate. Further studies using helium in cell culture or in clinical trials may clarify its clinical role for such diseases.

## Materials and Methods

The *in vitro* testing in this study was performed using a standard fluorometric furin activity assay (Anaspec: Fremont, CA). This assay uses a fluorometric substrate to measure the enzymatic activity of recombinant human furin. When the enzyme is active, it breaks the substrate in such a way that a chemical compound is released that will fluoresce when exposed to light at the proper wavelength. The assay includes appropriate quantities of the reagents for microplate studies and is designed for screening substances for furin inhibition as part of a medication discovery process.

After the reconstitution of the furin enzyme and the fluorogenic substrate, the reconstituted substrate was then placed in the control wells of the microplate (C1, C2, C3; D1, D2, D3; E1, E2, E3; and F1); and also in the test wells (C10, C11, C12; D10, D11, D12; E10, E11, E12; and F12). Subsequently, half (400 μl) of the furin enzyme solution was placed in a Schlenk tube, while the other half (400 μl) remained in its test tube. Schlenk tube are special glass tubes that have a side port for the addition or removal of gases. They are useful for studies involving a gaseous reagent or product.

At this time, the air was then gently removed from the Schlenk tube with a manual vacuum pump. None of the liquid containing the enzyme was visibly removed when the air was removed. The Schlenk tube was subsequently attached to a source of balloon-grade helium gas (Airgas), and helium gas was added to a pressure of approximately 5 psi above atmospheric pressure. After filling the flask to this pressure, the helium gas was disconnected and the hose to the Schlenk tube was clamped. At this time, the untreated furin was transferred from its test tube to the control wells of the microplate, and the helium-exposed furin enzyme in the Schlenk tube was poured into a separate test tube and then placed in the test wells. This additional step was taken because the 10 mL Schlenk tube was too long and narrow to permit the micropipette to access the 400 microliters of fluid at the distal end of the Schlenk tube. Unfortunately, this additional step likely increased our atmospheric helium losses from the Schlenk tube during the experiment and may contribute to a potential underestimate of the full inhibitory effect of helium on furin in this experiment.

The black, 96-well, non-binding surface microplate was subsequently placed in our Tecan Infinite F200 microplate reader (Tecan Group Ltd: Männedorf, Switzerland) and underwent automatic fluorescence measurements every 72.5 seconds for 24 hours. 72.5 seconds is the shortest time interval at which our microplate reader can provide continuous measurements of all 96 wells on the plate. The experiment could be optimzed in the future by limiting the microplate reading process to the control and test wells, rather than the entire microplate.

The mean fluorescence from the helium-exposed wells in series E compared to control are presented in Figure 1. Figure 2 includes the data in Figure 1, plus the data used to derive the mean values which are graphed in Figure 1 (the three individual sets of control and test wells). Figure 3 uses the data from Figure 2 to compare the ratio of the average furin activity with helium versus without helium over the 24 hour period of the experiment. An initial subset of the measured data is also presented in tabular form in Table 1.

There are two primary reasons why this paper emphasizes the findings from the E-series wells. First, these are the only samples for which our device obtained reliable measurements throughout the full 24 hour duration of the experiment. The C-series wells had such high levels of fluorescence after the first hour of measurement that the microplate reader stopped recording the measured fluorescence and merely recorded the error message “OVER”. The D-series control wells had a similar problem, but for a shorter duration of time. This limits our ability to analyze the data for these wells after the first hour, which is why the graphs and data presented from the C- and D-series wells only extends to the first hour of the reaction. This is perhaps not a great loss, as the fluorometric assay does not appear to have been validated for measuring duration of inhibition beyond 1 hour. The E-series wells did not have this excess fluorescence error, perhaps because they had a slightly smaller amount of furin enzyme present. Therefore, it is from the series E data that the tentative observation of a 16-hour duration of helium-based furin inhibition is drawn.

The second reason this paper emphasizes the findings in the E-series wells is that these samples, which were drawn and placed in the microplate last, are our best available demonstration of the degree to which helium is capable of inhibiting furin. This is consistent with our previous experiment with xenon as a furin inhibitor^22^, in which the xenon-treated furin samples which were drawn last from the tube and placed on the microplate last demonstrated the most effective inhibition. The cause of this trend is not clear; it occured with both xenon (heavier than air) and helium (lighter than air). Further study will be necessary to evaluate the cause of this trend.

(Of note, because of transfer losses, well F1 and well F12 did not receive sufficient enzyme to be considered reliable samples and were excluded from analysis.)

## Acknowledgements

The work of those who developed and produced the Anaspec furin activity assay made this study much easier than it otherwise would have been. Their service is gratefully appreciated. The assistance of the staff at Airgas is also recognized.

Dr. Beihl performed the research and authored this entire paper. He holds numerous patents pending on pharmacological compositions and methods of using noble gases to inhibit proprotein convertases, including methods of their clinical use in the treatment of cancers, certain viral and bacterial infections, degenerative diseases, and vascular diseases, as well as autoimmune, neurological, genetic, and other diseases. No governmental, institutional, or other external funding was received to perform this research.

## Appendix A

As shown in Figure 4, the inhibitory effect of helium is visible, even when the data from the Series C and D wells (which demonstrated less furin inhibition) is added to the Series E data. Figure 5 shows the previously mentioned trend of increasing inhibition from the Series C wells to the Series D and E wells. The inhibition in Series C alone is rather modest (Figures 6, 7, and 8), but in the Series D wells, increased inhibition is noted (Figures 9 and 10). The inhibition started at 55%, then decreased to 40% before increasing slowly toward 55% by the end of the hour (Figure 11). The slow increase in inhibition in Figure 11 may represent asymmetric enzymatic and substrate degeneration, rather than an increasing effect of helium over time. Further studies may clarify this subject.

**Figure 4.**
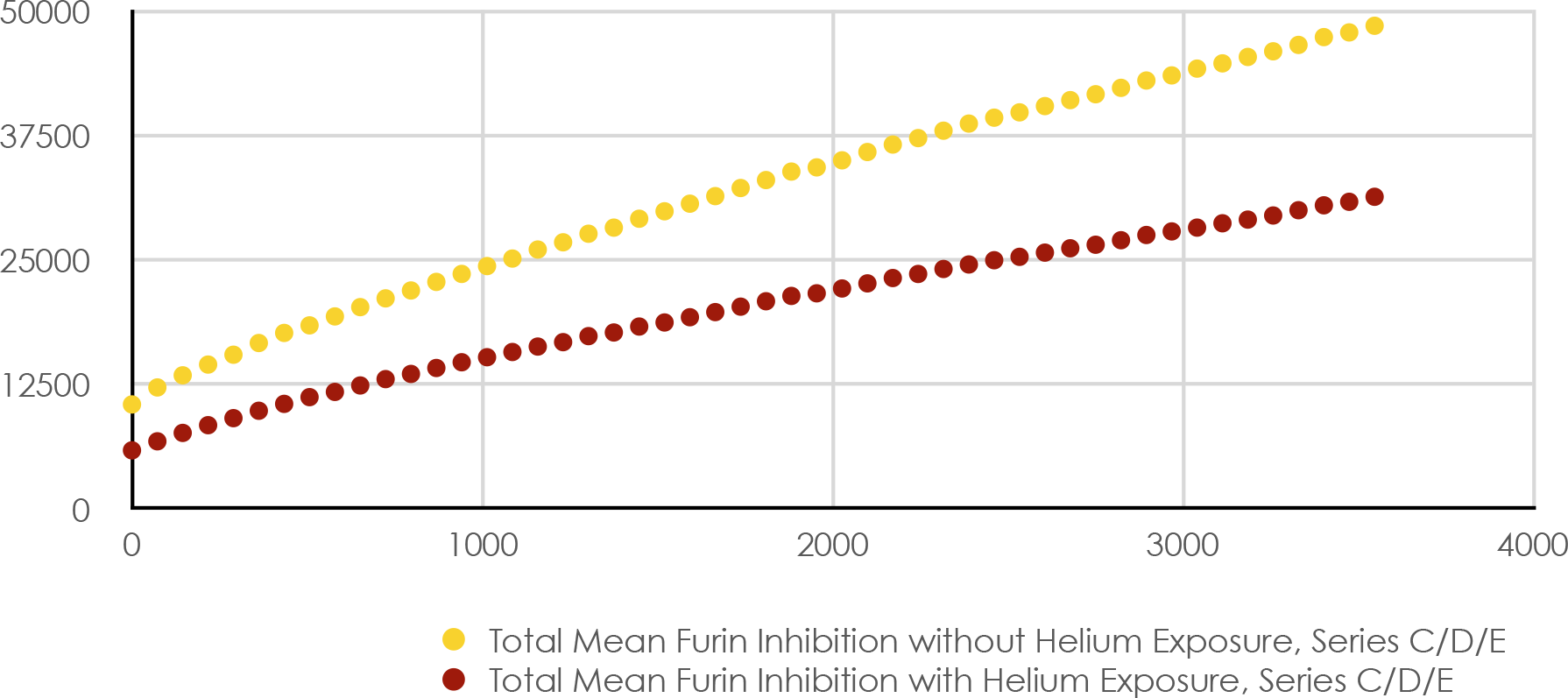
Mean Furin Activity with and without Helium Exposure vs. Time (s), Series C, D, E wells

**Figure 5.**
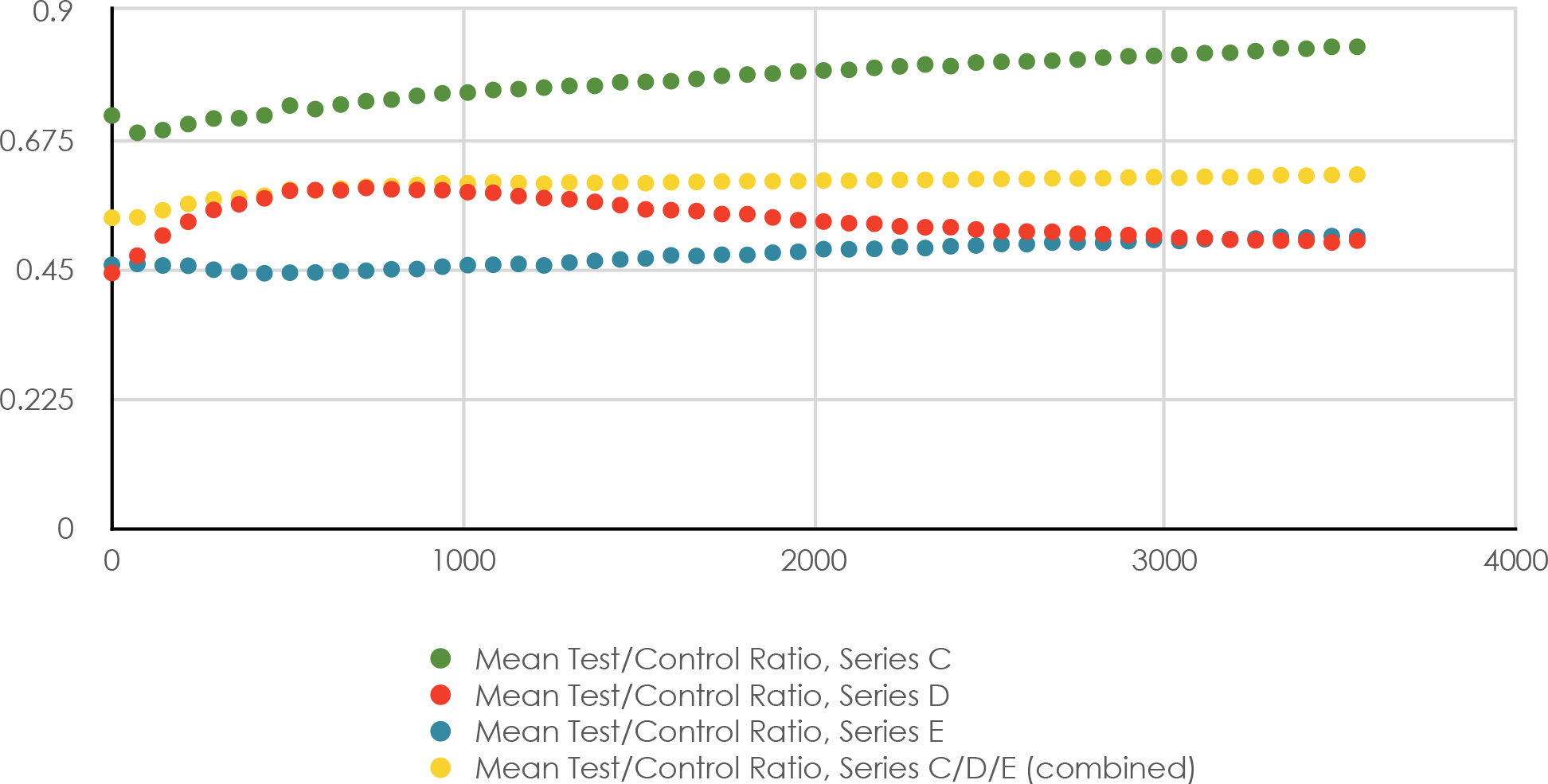
Mean Test/Control Furin Activity Ratio, Series C/D/E Wells

**Figure 6.**
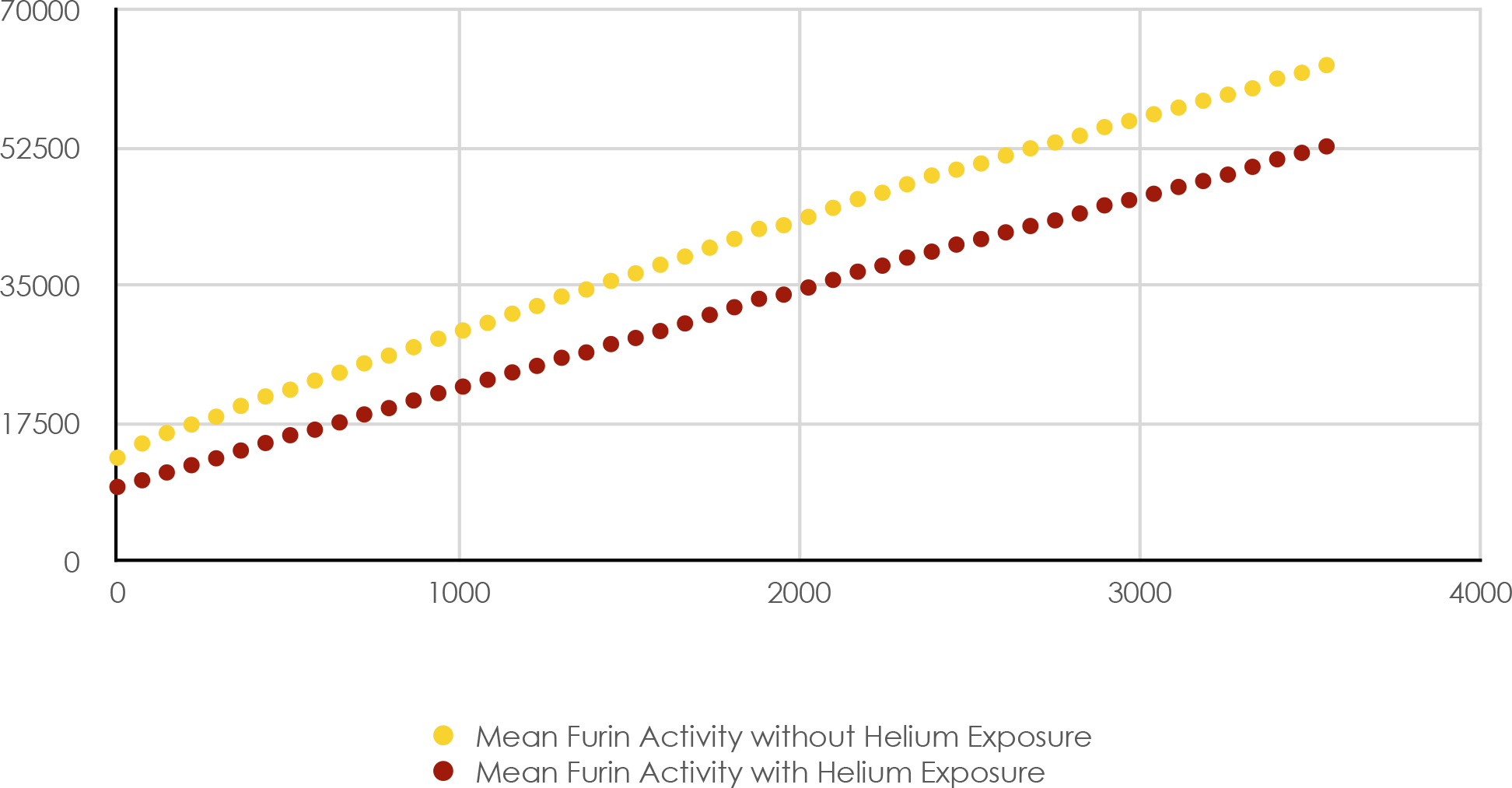
Mean Furin Activity with and without Helium Exposure vs Time (s), Series C Wells

**Figure 7.**
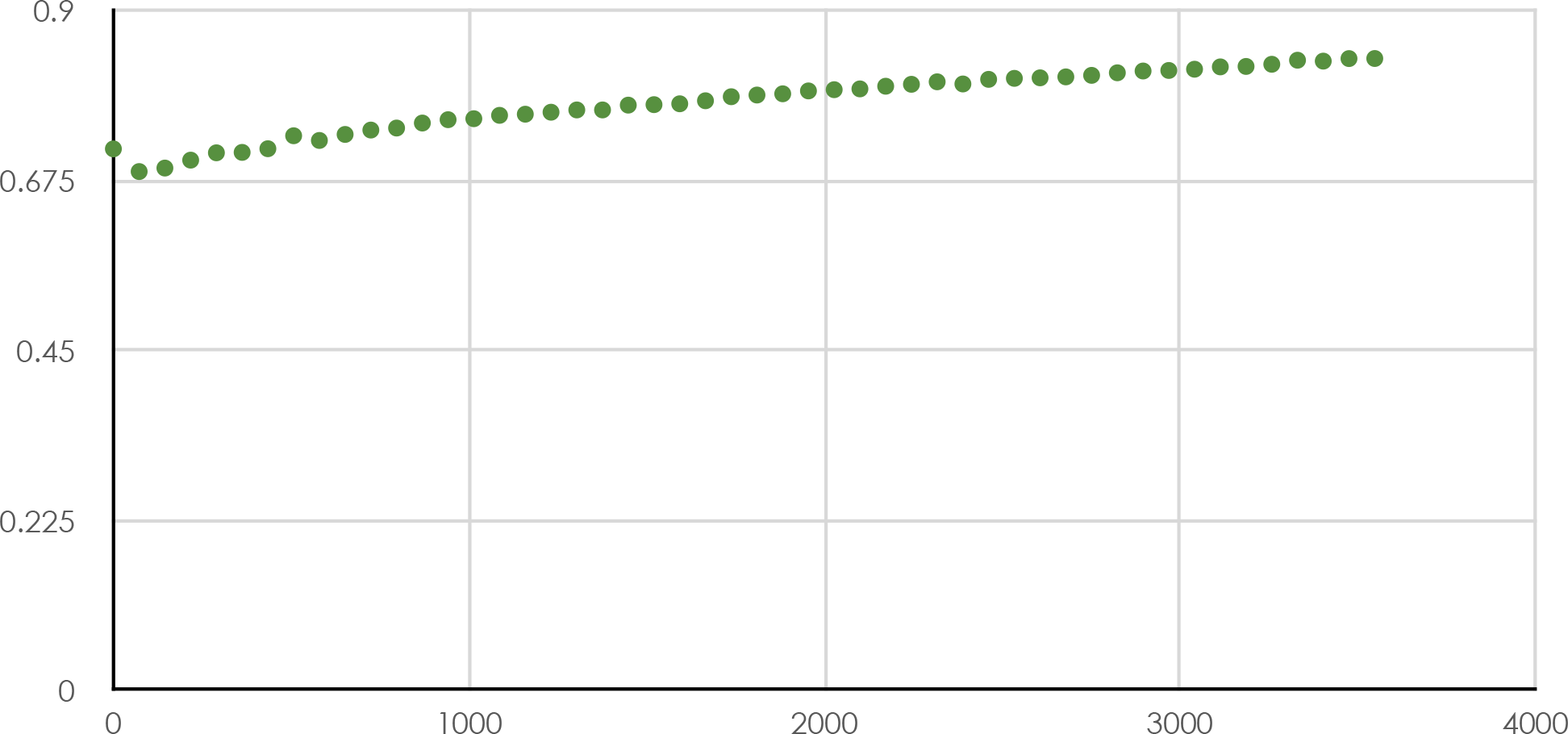
Mean Test/Control Furin Activity Ratio vs Time (s), Series C Wells

**Figure 8.**
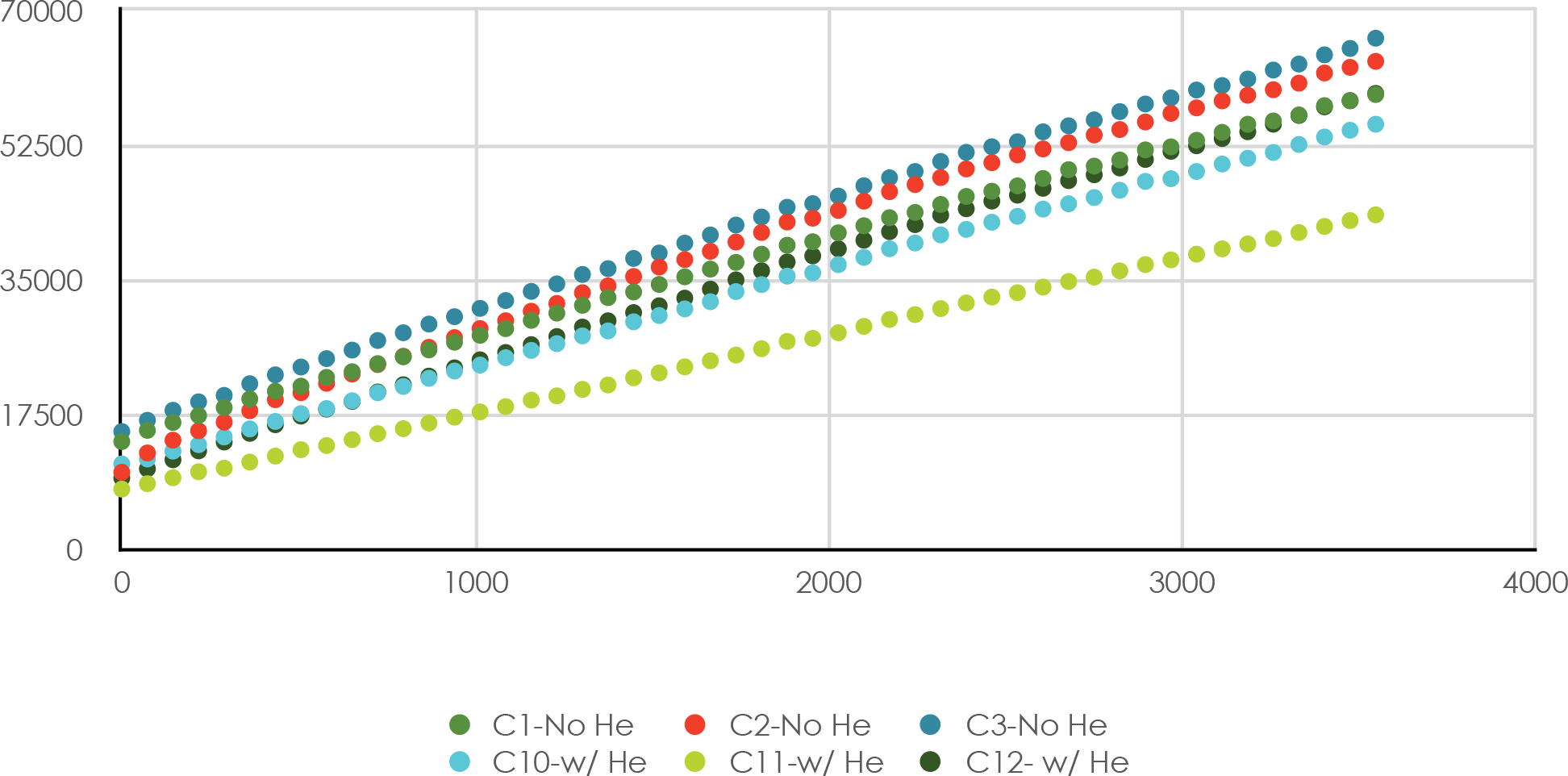
Furin Activity with and without Helium Exposure, vs Time (s), Series C Wells

**Figure 9.**
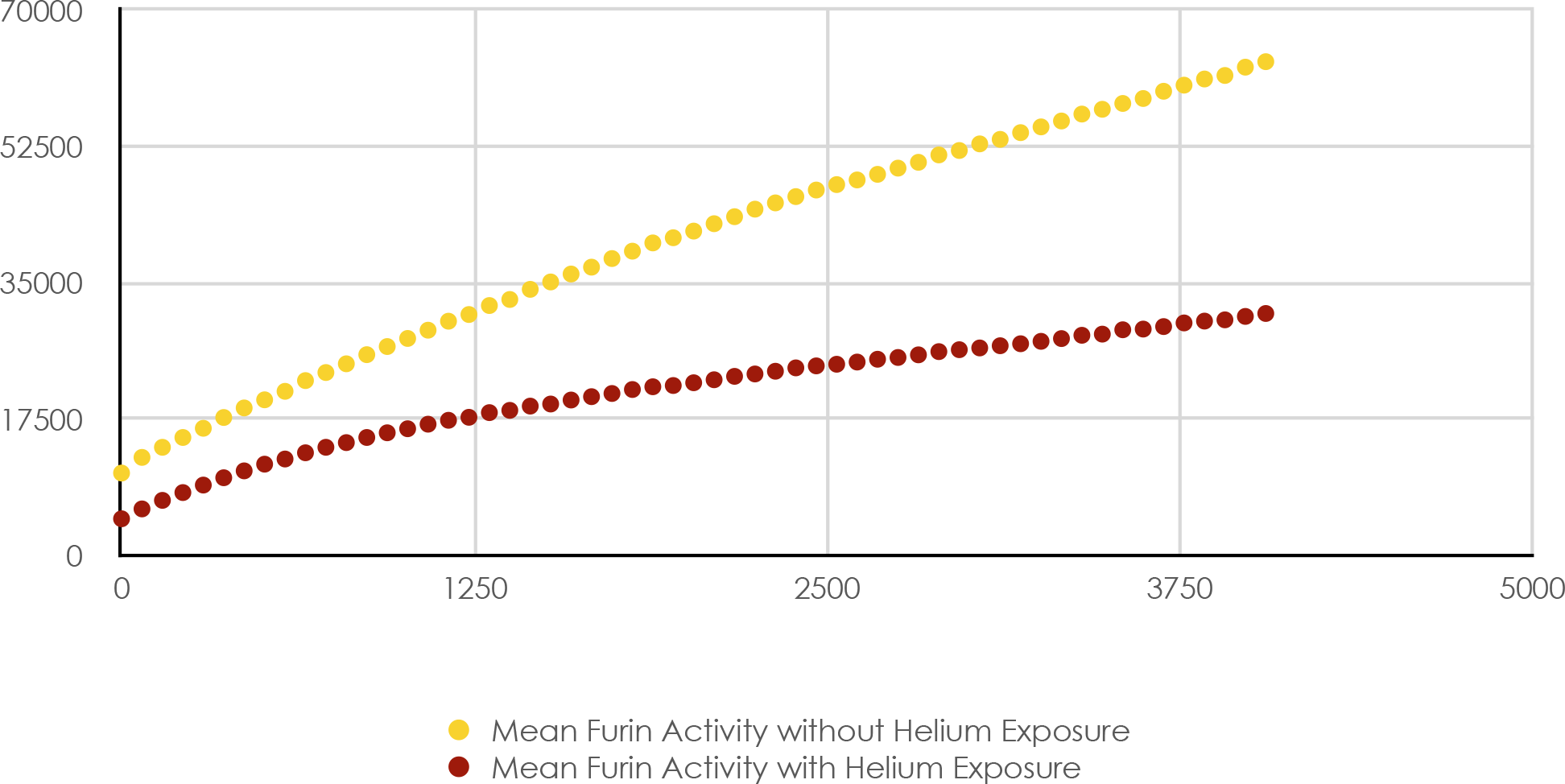
Mean Furin Activity with and without Helium Exposure vs Time (s), Series D Wells

**Figure 10.**
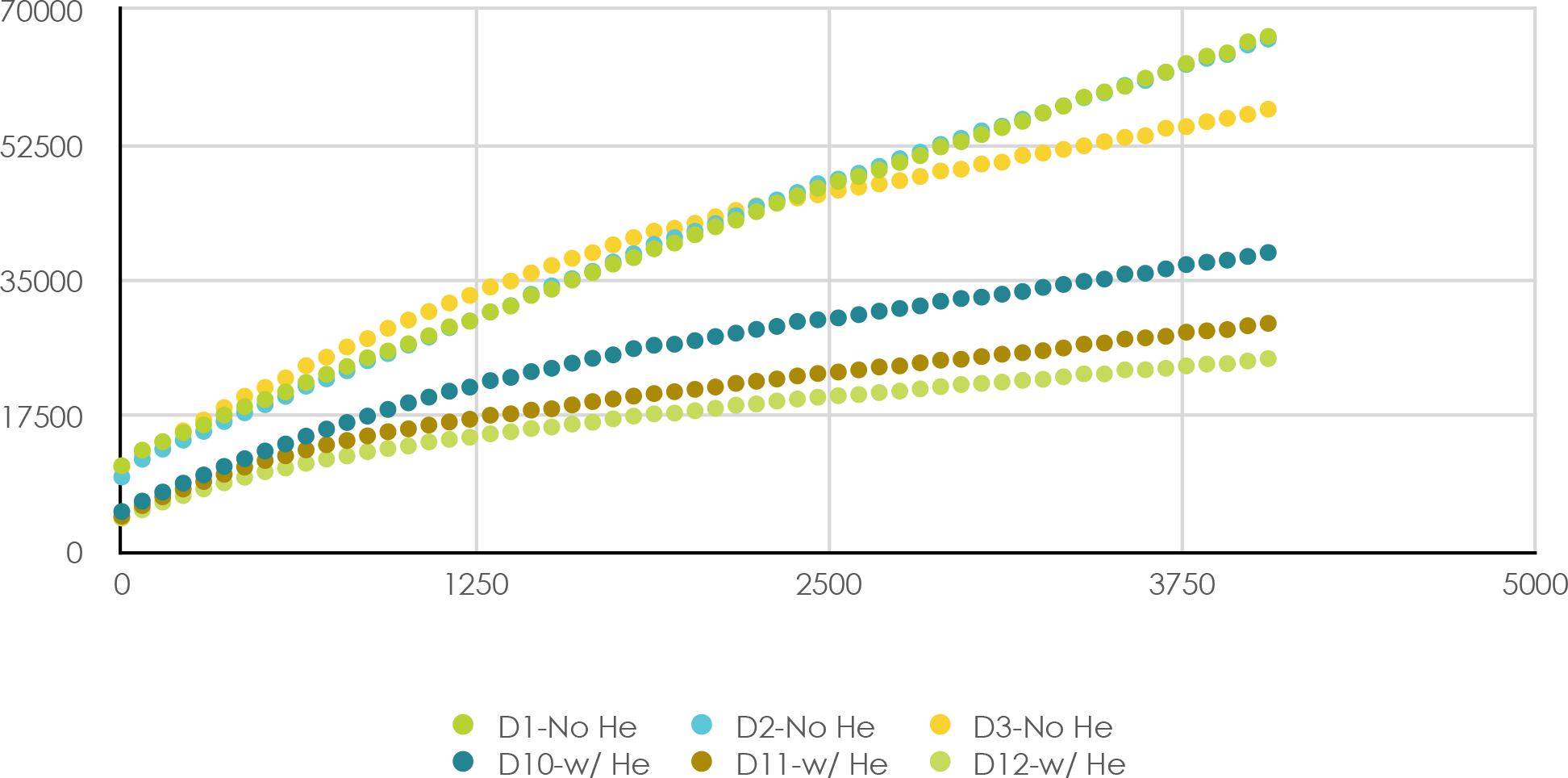
Furin Activity with and without Helium Exposure vs. Time (s), Series D Wells

**Figure 11.**
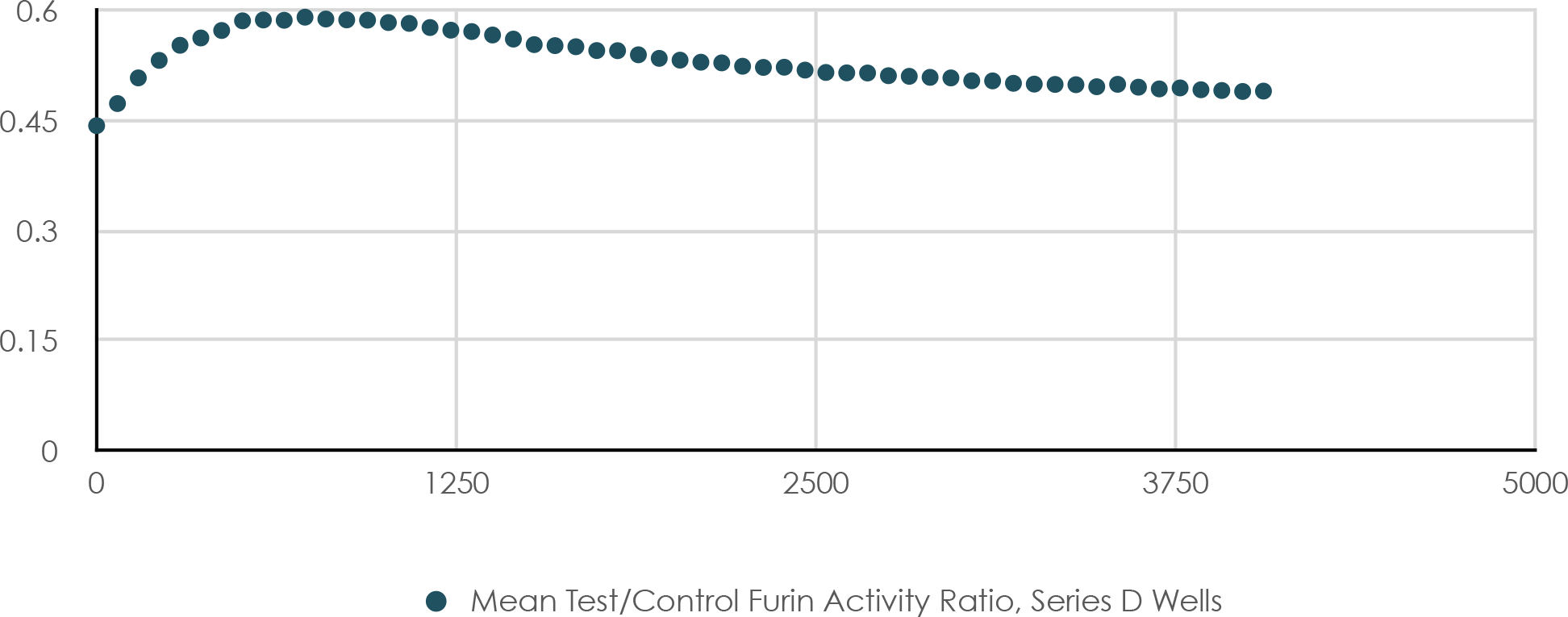
Furin Activity with and without Helium Exposure vs. Time (s), Series D Wells

